# Differential k-mer distributions enable identification of tetrapod subterminal heterochromatin

**DOI:** 10.64898/2026.10.19.760198

**Authors:** Robert Dalipovic, Adam M Session

**Affiliations:** Binghamton University

## Abstract

Subterminal heterochromatin are dynamic regions of the genome that are repeat rich and vary in structure between individuals of the same species. With advances in genome sequencing technology, these regions are being assembled for the first time in many groups such as *Xenopus*. We develop a k-mer based algorithm capable of identifying previously characterized subterminal satellites, as well as lower density satellites in diploid primates and *X. tropicalis*. When applying our algorithm to the allotetraploid genome of *Xenopus laevis*, we find that the subterminal heterochromatin regions include ancient subgenome-specific repeats that originated while the *X. laevis* progenitors were distinct species. This pattern challenges the model proposed in primates of ectopic recombination driving non-homologous chromosomes to share repeat sequences.

---

Subterminal heterochromatin (STH), or subtelomeres, are dynamic genomic regions of the chromosome found between the telomeres and the main chromosome body (interstitial region)^1^. Non-homologous chromosomes share satellite repeats and segmental duplications, which has been proposed to be maintained through non-homologous ectopic recombination ^2^. They are characterized in part by their high rates of recombination, structural mutations, and gene birth/death processes^1,3–5^. Furthermore, these regions are repeat rich making them difficult to assemble and distinguish orthologous between paralogous sequences. Thus, these regions were initially only studied and defined in humans and other primates as well as more widely used models such as *Saccharomyces cerevisiae* and *Plasmodium falciparum* ^1,3,4,6–8^. While subtelomeres are presumably present in all eukaryotic genomes, there is no similarity in size or repeat content between distantly related species like primates, yeast, and *Plasmodium*^1^.

Increased availability of long-read sequencing technologies has enabled more accurate assembly of highly repetitive regions. This has led to characterization of STH in more systems such as gorilla, chimpanzee, and the diploid amphibian *Xenopus tropicalis* ^5,9–11^. *X. tropicalis* STH regions are around 20 megabases(Mbp) in length, in contrast to primate regions which average ∼5 Mbp^5,9,11^. STH structure in *X. tropicalis* includes shared repeats between non-homologous chromosomes, similar to primates. The identification of these regions in *X. tropicalis* is based on the increased recombination rate in the STH compared to the interstitial regions. STH repeats are classified as satellites and retrotransposons, the most common being a 52-base pair(bp) satellite^5^.

Polyploidy, or whole genome duplication, has long been recognized as a source for genomic novelty through providing raw material for evolutionary diversification^12,13^. Polyploidy occurs through two primary mechanisms: autopolyploidy (genome duplication within one species) and allopolyploidy (genome duplication through merging the diploid complements of two distinct diploid species)^14–16^. Allopolyploids are distinct because their genomes will show mostly bivalent inheritance at the time of hybridization, unlike autopolyploids which have a period of multivalent inheritance. Recent advances in long-read genome sequencing technology and methods have made the sequencing and assembly of chromosome-scaled polyploid genomes easier. Despite these advancements, analysis of polyploid genomes has largely been restricted to understanding how the genomes respond to the increased gene/chromosome copy number, classifying subgenomes of allopolyploids without extant progenitors through asymmetric repeat densities, and how the subgenomes of allopolyploids show asymmetric evolution (biased fractionation)^17–21^. Given the unique natural experiment provided by allopolyploidy (merging two distinct diploid complements into a single species), allopolyploid genomes are currently an underutilized tool for studying eukaryote genome evolution.

Here, we present a novel computational method to identify tetrapod subterminal heterochromatin regions using diagnostic k-mers, ANOVA, and a Hidden Markov Model. We show that this method identifies previously characterized regions and sequences in diploid primates and *Xenopus tropicalis* before applying the algorithm allotetraploid *Xenopus laevis*. For the diploid genomes with previously characterized subterminal satellites, our method identifies not only the previously characterized sequences, but additional satellites with increased subterminal density. Additionally, the co-localization of the subterminal heterochromatin with subgenome-specific repeat families in allotetraploid *X. laevis* challenges the existing model of non-homologous recombination as the expansion mechanism for subterminal heterochromatin in tetrapods.

## Results

### Generalized method for K-mer based identification of subterminal heterochromatin

We have previously used asymmetric k-mer distributions to identify families of repeats that differentiate subgenomes of allopolyploids ^20,21^. Briefly, normalized k-mer counts for each chromosome are used to look for k-mers that differentiate homoeologous chromosomes (those related through the genome duplication event) within each chromosome set. Taking a union of k-mers across all chromosome sets will reveal what k-mers differentiate subgenomes which can then be used to identify specific repeat families based on co-localization in the genome. ANOVA is used to assess how well each k-mer is able to differentiate between the chromosome sets.

The problem of subterminal heterochromatin identification is similar, differential repeats in the subterminal regions of chromosomes should leave a clear signal in the normalized k-mer density compared to the interstitial regions. This analysis is outlined in Figure 1. The initial step is trimming the ends of each chromosome to generate subterminal and interstitial regions. This requires knowledge of the rough expected subterminal size in the species, 5Mbp for apes, and 20Mbp for amphibians. Once trimmed, normalized k-mer counts (k-mer count/bp) are computed for each segment, and an ANOVA is done to look for k-mers that differentiate the interstitial regions from the subterminal regions of non-acrocentric chromosomes. In the case of allopolyploid *X. laevis*, ANOVA categories also consider subgenome (i.e. L-interstitial, L-subterminal, S-interstitial, S-subterminal; Figure 1b).

**Figure 1.**
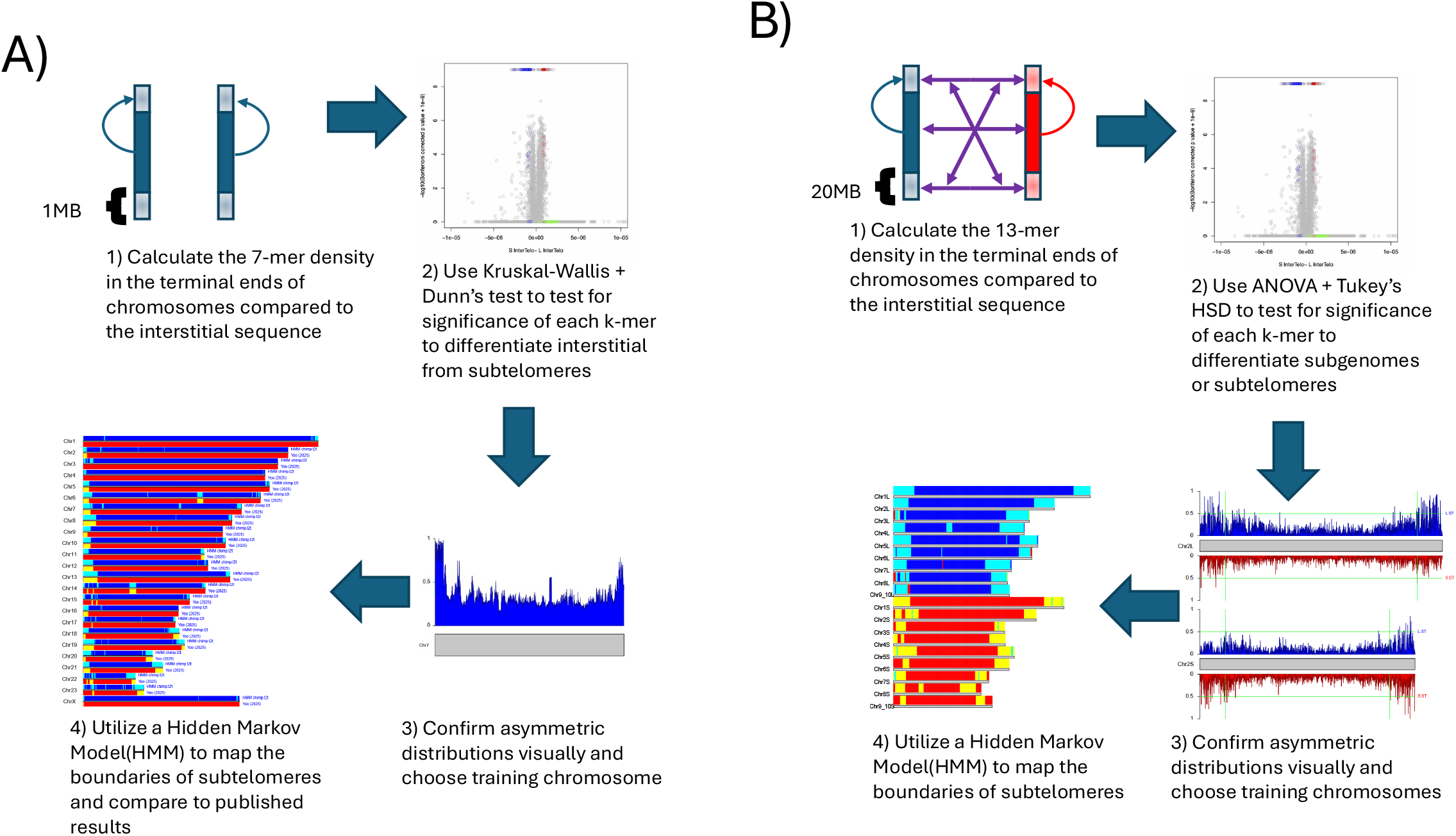
Outlining strategies to identify subterminal heterochromatin enriched k-mers a) Strategy to identify subterminal k-mers in diploid ape genomes (*P. troglodytes, G. gorilla*). (Top left) First, the terminal 1MB of each chromosome is removed and analyzed separately, and 7-mer density is calculated for each segment. (Top right) Next a Kruskal-Wallis and Dunn’s test are used to assess the ability of each 7-mer to differentiate terminal vs interstitial segments. (Bottom right) These asymmetrical distributions are confirmed on each chromosome, and the best looking is chosen as the training chromosome for the HMM. (Bottom left) Finally the HMM is used to map the boundaries based on the 7-mer density in 100-kb bins. This is visualized and compared to previously published results. The diploid *X. tropicalis* was done similarly, except for k=13 and longer 20Mbp terminal regions. b) A similar strategy for the allotetraploid *Xenopus laevis* which incorporates subgenome identity as well as terminal vs interstitial for the chromosomes. There must be one training chromosome for each subgenome of the allotetraploid.

The size of the subterminal regions will also dictate the k-mer length used. For the longer subterminal regions of *Xenopus*, we use k=13 the same as was used for chromosome-wide subgenome-specific repeats. For the smaller regions of primates, k=13 is unlikely to appear by chance in a 5 million bp region, so we lower to k=7. 7-mers have a bimodal genomic distribution that requires the non-parametric Kruskal-Wallis and Dunn’s test^22^ to assess significance for each 7-mer to differentiate the genomic regions. A complete discussion of the impact of *k* length on identifying differentially distributed repeats is included in Supplemental Note 1.

Once k-mers that are significantly enriched in the terminal regions are identified, they can be mapped back to the genome in order to confirm their asymmetry, as well as refine exact boundaries for each subterminal region. We do this by utilizing a Hidden Markov Model (HMM) that looks at the density of subterminal k-mers in each 100-kilobase (kb) bin to map fine boundaries to all chromosomes, including the acrocentrics not used in the initial step of the analysis. The mapping of the k-mers to the genome also allows us to extend the smaller k-mers into larger satellite consensi as well as look for overlapping satellite repeats that can inform how the specific satellite may have evolved.

We have applied this algorithm to 4 tetrapods: the primates chimpanzee (*Pan troglodytes*), Gorilla (*Gorilla gorilla*), as well as the amphibian models *Xenopus tropicalis* and *Xenopus laevis*. The ape genomes were chosen for their deep characterization of the pCht satellite with respect to subterminal heterochromatin ^9,10^. The genome of *Homo sapiens* was not included primarily due to the absence of the pCht satellite as a positive control, as well as the common practice of removing the dynamic subterminal regions of human chromosomes onto unlocalized scaffolds. *X. tropicalis* also has defined subtelomeric regions and satellites^5^. *X. laevis* was chosen as the closest allopolyploid system for comparative analysis with primates to test the model that ectopic recombination is the mechanism used to spread similar repeats into non-homologous chromosomes.

### K-mer based identification of ape subterminal heterochromatin

For chimpanzee and gorilla, we tested our algorithm on the T2T genome assemblies (Supplementary Note 2). For both primate T2T assemblies we use a single haplotype, however in Chimpanzee we use the “Golden Path” haplotype which a mix of paternal and maternal assemblies. The primary subterminal satellite for chimpanzee and gorilla (pCht or StSat) was identified previously by telomere amplification^23^. For both species, the subterminal regions were recently refined based on mapping of the pCht satellite via BLAST ^9^. If our algorithm is correctly defining the satellites of subterminal regions, then we should identify the pCht satellite in our analysis, as well as any other satellites that are accumulating in these regions that have not been characterized. In chimpanzee, our algorithm identifies 207 7-mers that are significantly higher in the subterminal region than interstitial. Of these 7-mers, 31 overlap with pCht across at least 6bp, while 140 overlap with 4bp or more of pCht. For gorilla, 496 7-mer are enriched in the subterminal regions,196 of which overlap with pCht across 6bp or more, while 434 overlap at 4bp or more. In both species, all positions of pCht overlap with subterminal 7-mers (Fig. 2a). The high number of 7-mers in gorilla overlapping with the 32-bp repeat is due the larger number of alleles segregating in gorilla subterminal heterochromatin compared to chimpanzee ^9^. Overall, these data suggest that our method is able to identify pCht independently of the methods that have been used previously.

**Figure 2.**
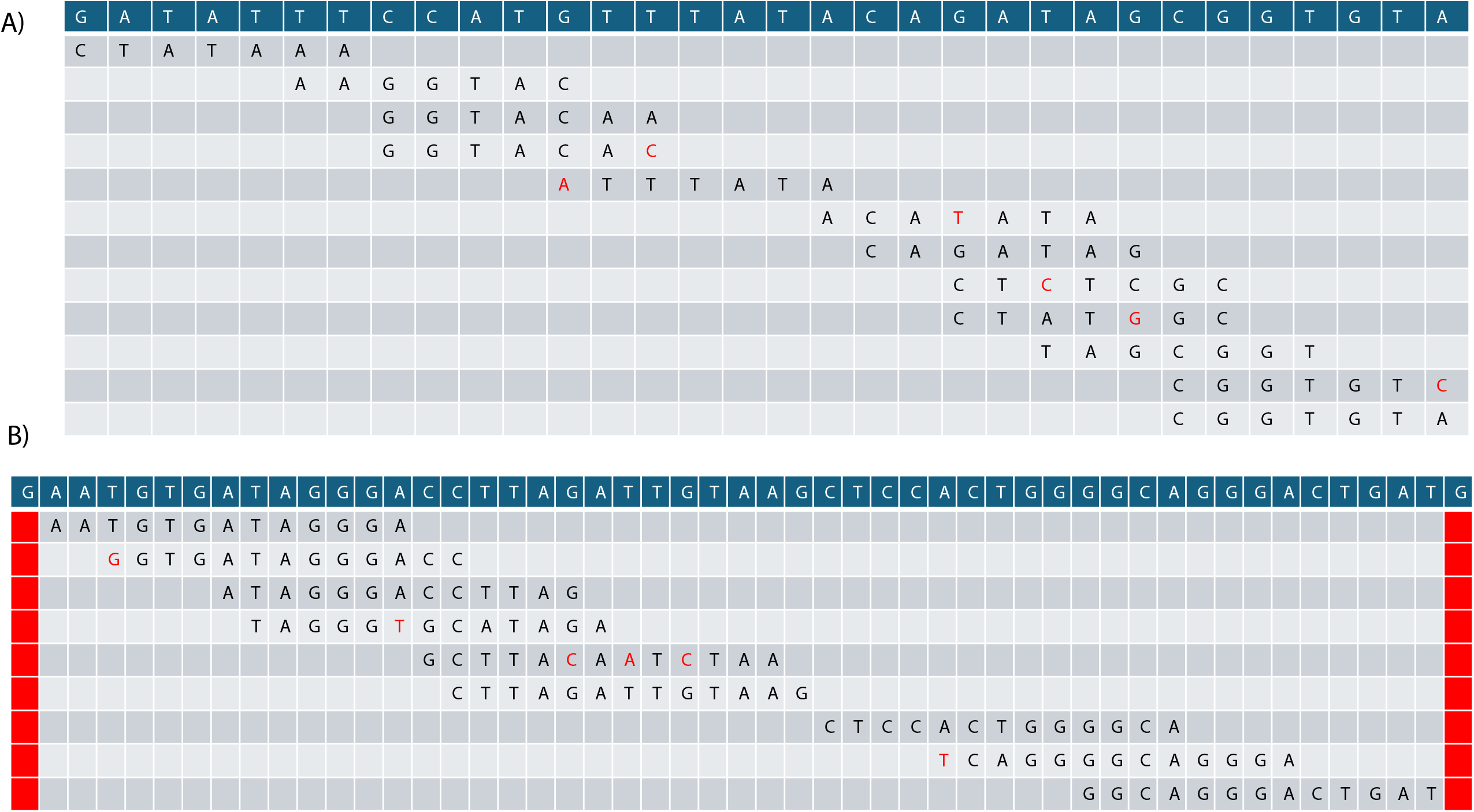
Alignment of subterminal k-mers against previously characterized subterminal satellites in primates and *X. tropicalis* a) Top row is the pCht consensus identified by Royle et al. Subsequent rows are a subset of the 7-mers that we find to be subterminal that align to pCht. Red letters indicate SNPs in the 7-mer compared to the consensus. Rows with asterisks indicate 7-mers where the ‘canonical’ sequence used by Jellyfish is the reverse complement of the consensus. b) Top row is the 52-mer consensus identified by Bredeson et al. Subsequent rows are a subset of the subterminal 13-mers that align to the 52-mer consensus. Red letters indicate SNPs in the 13-mer compared to the consensus. Red columns indicate positions in the consensus that have no strict alignments (1bp on either terminal end of the consensus)

In order to identify the specific repeat families our differential k-mers represent in primates we first overlapped our k-mer loci with RepeatMasker annotations. While there was significant overlap, many of the repeat families were given generic names (e.g rnd-1-family-436). In order to assess if we are able to identify pCht as a subterminal satellite without access to the published consensus, we instead ran Tandem Repeat Finder^24^ (TRF) on each genome to first call satellite arrays, then ask if our subterminal k-mers overlap with these consensi. Using this method, we identify pCht satellite arrays as subterminal, as well as a number of non-pCht satellite arrays (Supplemental Note 2)

The HMM in primates measures the amount of subterminal 7-mer density ‘emitted’ from each 100-kb bin in order to assign each bin as interstitial or subterminal based on a training set (Figure 3). For both chimpanzee and gorilla, Chr7 was used based on having a good pattern through visual inspection (Supplemental Figure 3). We predict a total of 337.9Mbp and 584.1Mbp of subterminal sequence in chimpanzee and gorilla respectively compared to a total of 159.5Mbp and 291.1Mbp predicted by Yoo^9^.

**Figure 3:**
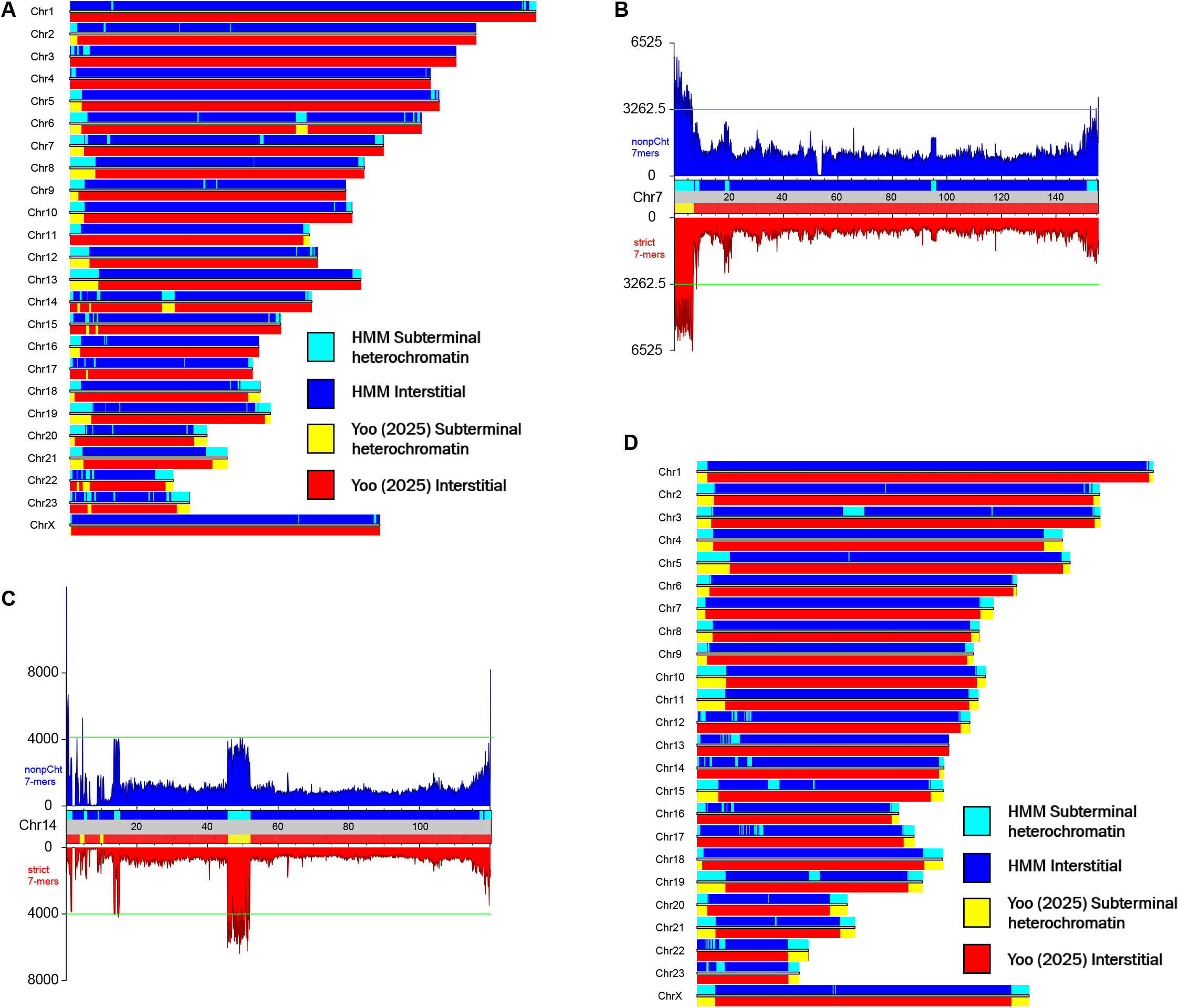
Primate subterminal heterochromatin loci comparison with previous results A) Karyogram comparing predictions of chimpanzee subterminal heterochromatin regions between our HMM method and Yoo et al 2025. Our prediction is shown in blue/light blue above the chromosome, while theirs are shown in red/yellow below the chromosome. B) Karyogram of chimpanzee Chromosome 7, showing the HMM/Yoo identity from A) in the center, and the density of non-pCht 7-mers (top;blue) vs strict(6/7 matches) pCht 7-mers (bottom; red). At the 3’ end of the chromosome, we predict a subterminal region that Yoo does not, and the non-pCht 7-mers seem to be driving this prediction moreso than pCht 7-mers. C) Karyogram of chimpanzee Chromosome 14, showing the HMM/Yoo identity from A) in the center, and the density of non-pCht 7-mers (top;blue) vs strict pCht 7-mers (bottom; red). The y-axis maximum was manually set to 8,000 in order to better visualize the differences, despite >8,000bp of coverage in the first bin for non-pCht. At the 5’ end of the chromosome the two methods disagree on the exact position of the subterminal heterochromatin, driven largely by differences in the pCht vs non-pCht 7-mer distribution. Again, the 3’ end of the chromosome, we predict a subterminal region that Yoo does not, and the non-pCht 7-mers seem to be driving this prediction moreso than pCht 7-mers. D) Karyogram comparing predictions of gorilla subterminal heterochromatin regions between our HMM method and Yoo et al 2025. Our prediction is shown in blue/light blue above the chromosome, while theirs are shown in red/yellow below the chromosome.

In chimpanzee, we call subterminal regions for 47/48 chromosome ends in this haplotype, compared to 27/48 identified by Yoo et al using only pCht (Figure 3a; Supplemental Note 2). We identify both blocks of interstitial subterminal satellites identified by Yoo et al. on chromosomes 6 and 14 in the chimpanzee genome. There are 25/27 chromosome arms with subterminal predictions by both methods. 27/50 overlapping borders (5’ and 3’ of the 25 overlapped regions) differ by <100,000 bp, which given the difference in methodology is as close as our predictions could be. In the remaining overlapping segments, our k-mer based algorithm often predicts longer subterminal regions, likely due to taking into account more satellites than just pCht. Indeed, when splitting the k-mers into groups that overlap pCht and do not, we find that the regions exclusive to our methods are due to the non-pCht satellites (Figure 3b). There are two chromosome arms where our predictions do not agree (Figure 3c; Chr14). Our predictions in these regions are driven primarily by non-pCht sequences which are more sparse here than other chromosomes, which may explain the divergence. These are also acrocentric chromosomes which may not have contiguous subterminal regions in the smaller p arms.

For gorilla, we call subterminal regions for 47/48 chromosome ends, compared to 40/48 identified by Yoo et al using pCht (Figure 3d). We do have subterminal predictions on the p arm of acrocentric chromosomes, but these are smaller non-contiguous regions. We also identify multiple interstitial regions with a high concentration of subterminal repeats that were not identified by Yoo et al (Figure 3d). Our predictions overlap with Yoo et al. in 39/40 regions, and have a difference in border prediction of < 100kbp in 73/78 overlapping borders. The large overlap in regions predicted by independent methods is likely driven by pCht being higher density in the Gorilla genome, which gives that family of satellites larger weight in our algorithm and more similarity with the pCht mapping method in Gorilla compared to Chimpanzee.

Overall, our algorithm recovering pCht and novel subterminal satellites in ape assemblies is strong evidence that this method could work in species whose subterminal satellites are not as well defined.

### K-mer based identification of *Xenopus* subterminal heterochromatin

For *X. tropicalis*, we used k=13 due to the longer mean subterminal length (20Mbp) and wanting to compare *Xenopus* sequences to the 13-mers previously used to identify subgenome-enriched repeats (Supplemental Note 1) ^20^. We compared our *X. tropicalis* subterminal heterochromatin calls to those previously published by Bredeson et al on the long-read v10 genome assembly (Supplemental Note 3). Unlike primates, both methods predict subterminal heterochromatin on all chromosomes. For 9/10 chromosomes, we predict similar subterminal ranges to Bredeson et al, with the primary difference being our regions do not always extend to the end of the chromosome while the Bredeson prediction must extend to the end(Figure 4a). Our ranges are not consistently longer for *X. tropicalis* likely due to the difference in methodology: the previous predictions of *X. tropicalis* subterminal heterochromatin used segmental duplications to identify these regions, so it is not the case that we are capturing more information as described above for primates. For *X. tropicalis* Chr10, we predict much longer subterminal regions than were identified using segmental duplications. This may be due to the higher GC% of *X. tropicalis* Chr10, which has been shown to disrupt sequence-based genomic comparisons ^25^.

**Figure 4:**
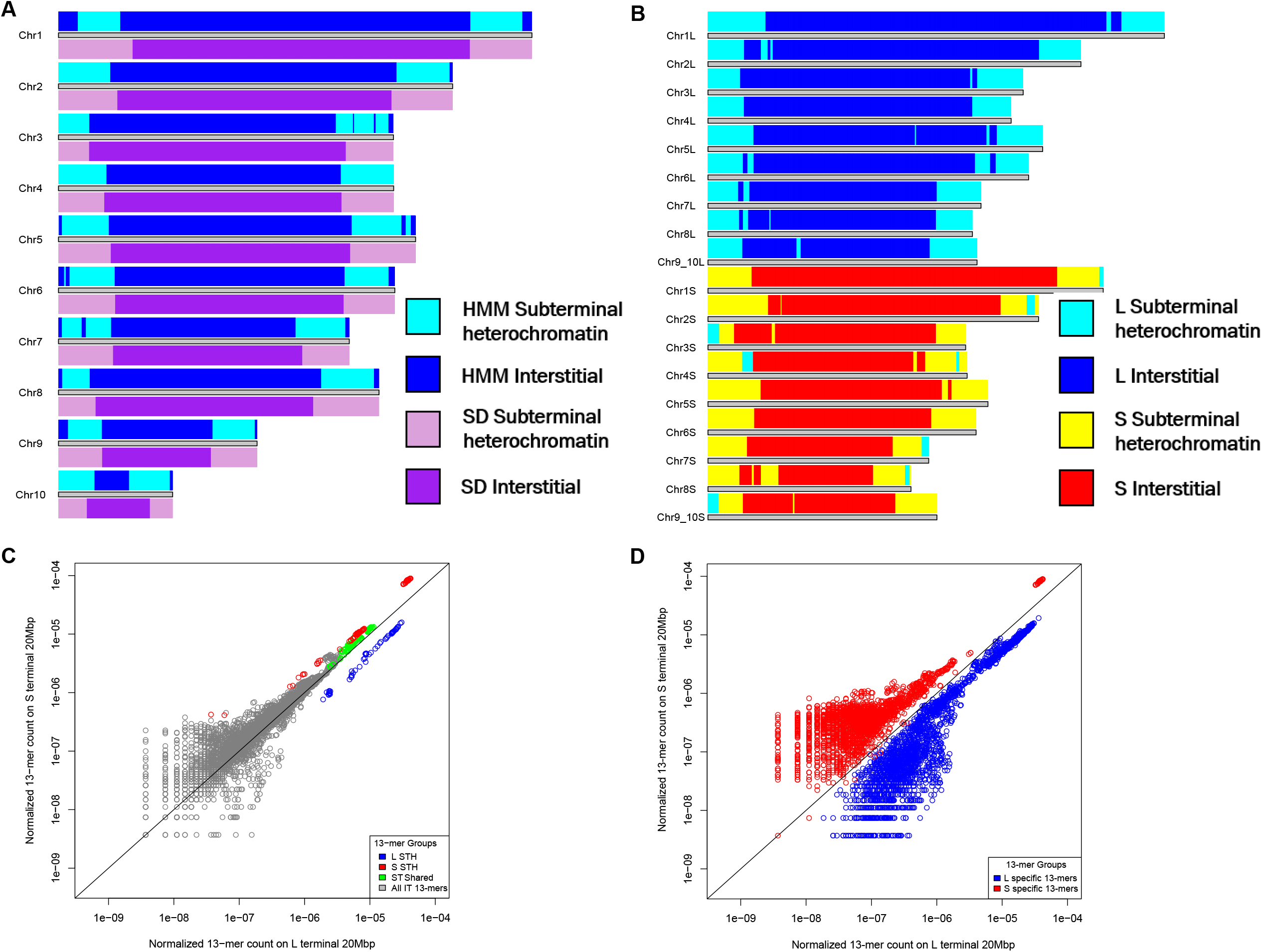
*Xenopus* subterminal heterochromatin loci A) Karyogram comparing predictions of *X. tropicalis* subterminal heterochromatin regions between our HMM method and Bredeson et al 2024. SD refers to the segmental duplication method used by Bredeson et al. Our prediction is shown in blue/light blue above the chromosome, while theirs are shown in purple/light purple below the chromosome. B) Karyogram showing HMM predictions in allotetraploid *X. laevis*. The HMM prediction is shown above each chromosome, blue/light blue for the L subgenome predictions, red/yellow for the S subgenome predictions. Some subterminal L regions are predicted on S chromosomes. C)Scatterplot comparing the mean normalized subterminal 13-mer count for the terminal 20Mbp on L chromosomes on the x-axis, and the terminal 20Mbp of S chromosomes on the y-axis. Black line is x=y. Embedded key shows color identity of each k-mer group. While the ANOVA does split L-subterminal 13-mers from S-subterminal 13-mers and both from those that are shared, it is important to note that all subterminal 13-mers have increased density on both subgenomes, just that some are higher on one subgenome vs the other. D)Scatterplot comparing the mean normalized subgenome-enriched 13-mer count for the terminal 20 Mbp of L chromosomes on the x-axis and the terminal 20Mbp of S chromosomes on the y-axis. Black line is x=y. The subgenome 13-mers still show a clear differential distribution in these genomic regions.

In *X. tropicalis*, we identify 1,961 13-mers, 25 of which overlap with the 52-mer identified by Bredeson et al. by 12 or more bp. This strict overlap recovers positions 2-51 of the 52-bp consensus, however the remaining 2bp are covered by other 13-mers that overlap the terminal end of the satellite and more flanking sequence (Figure 2b). We were able to identify the previously characterized 52-mer independently using TRF as well, although in TRF the consensus for most instances is actually a 53-mer that is extended by a single base pair..

Finally, we used our algorithm to identify the subterminal regions of allotetraploid *X. laevis* chromosomes. We identified 13-mers enriched in each of the 4 categories of the *X. laevis* genome (Figure 1b; L-subterminal, L-interstitial, S-subterminal, S-interstitial). For this HMM, the emission from each bin used not only the subterminal 13-mers, but also the subgenome-enriched 13-mers that differentiate the subgenomes (Supplemental Figure 5; Figure 4b; Supplemental Note 4). Many of the subterminal 13-mers are significantly enriched on one subgenome over the other, however unlike the ancient subgenome-specific repeats the subgenome-enriched subterminal repeats are shared between both subgenomes, simply having a higher density in one compared to the other (Figure 4c). The subterminal enriched sequences show no direct overlap with the subgenome-enriched repeats. Interestingly the subgenome-enriched repeats that represent the ancient transposable elements that expanded while the progenitor species were separated are still partitioned in the subterminal regions of *X. laevis* chromosomes (Figure 4d). This is relevant because previous analyses have hypothesized that the subterminal regions of chromosomes harbor related repetitive sequences due to non-homologous recombination^2,9,26^. If that were true for *X. laevis* subterminal heterochromatin, we would expect the ancient signal to be scrambled in these regions, which it is not. Therefore, we can conclude that despite the similarity in shared non-homologous duplications and repeats, *Xenopus* subtelomeres cannot be propagated through ectopic recombination.

**Figure 5:**
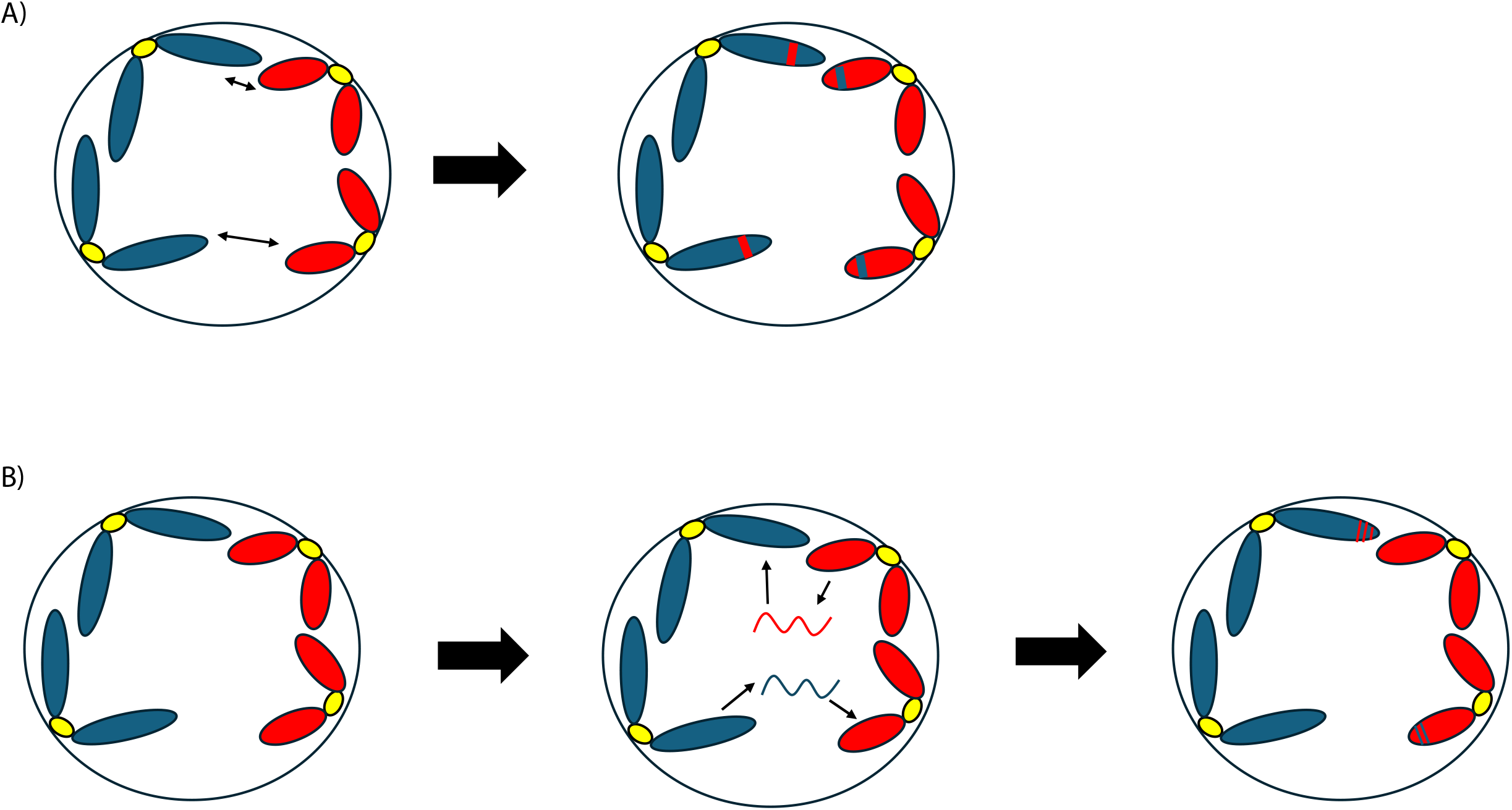
Model of subterminal heterochromatin expansion in allopolyploid genomes A) Illustration of how subterminal heterochromatin spread through ectopic recombination would appear in allopolyploid genomes. Here, the blue and red chromosomes are different subgenomes which have divergent families of transposons that ‘paint’ them distinctly. If subterminal heterochromatin was spread through ectopic recombination, we would expect to find small blocks of opposite subgenome identity in the newly assembled subterminal heterochromatin B) Illustration of our updated model of subterminal heterochromatin spread. Specifically, we hypothesize that the terminal ends of chromosomes come into close proximity within the cell, which allows satellite sequences to transpose between them without recombination. This model is consistent with results from both primates and *Xenopus*, and may apply to all eukaryotic genomes.

## Discussion

Our results show that asymmetric distribution of k-mers in the terminal ends of chromosomes can be used to rapidly identify satellite repeats that accumulate in the subterminal heterochromatin. The goal of many genome assemblies currently is to achieve a telomere-to-telomere or gapless assembly, however correct assembly and annotation of subterminal heterochromatin remains a challenge despite advancements in sequencing chemistry and assembly algorithms^11^. This method allows for rapid identification of these sequences even in gapped long-read assemblies such as *Xenopus*, which could prove useful to the correct assembly of subterminal regions in many species. While the subterminal satellites can be identified through methods like telomeric amplification^23^, it is currently more cost effective in non-model systems to have an algorithm capable of identifying the same sequences from the genome assembly alone.

The non-homologous ectopic recombination model of STH evolution in diploid genomes predicts that the subgenome-specific repeats of polyploids should be mixed in the terminal ends of allopolyploid chromosomes due to the non-homologous recombination that would be predicted after the hybridization of progenitor genomes into a single nucleus (Figure 5a). We find that the ancient subgenome-enriched repeats still partition L and S chromosomes in the subterminal regions, rejecting the hypothesis that these sequences could be expanding through non-homologous recombination in *X. laevis*. The fact that all subterminal signals are shared, even if they are biased by subgenome, still argues for non-homologous sequences sharing these satellite expansions. Our model is that these sequences spread through non-homologous transposition, perhaps mediated by the unique satellite expansions for each species (Figure 5b).

In summary, our study advances our ability to identify subterminal heterochromatin as well as our understanding of the biology of these complex genomic regions. Additionally, we illustrate how chromosome-scaled allopolyploid genomes are unique natural experiments in genome evolution that are underutilized for answering questions that apply to diploid genomes as well.

**Supplemental Figure 1:**
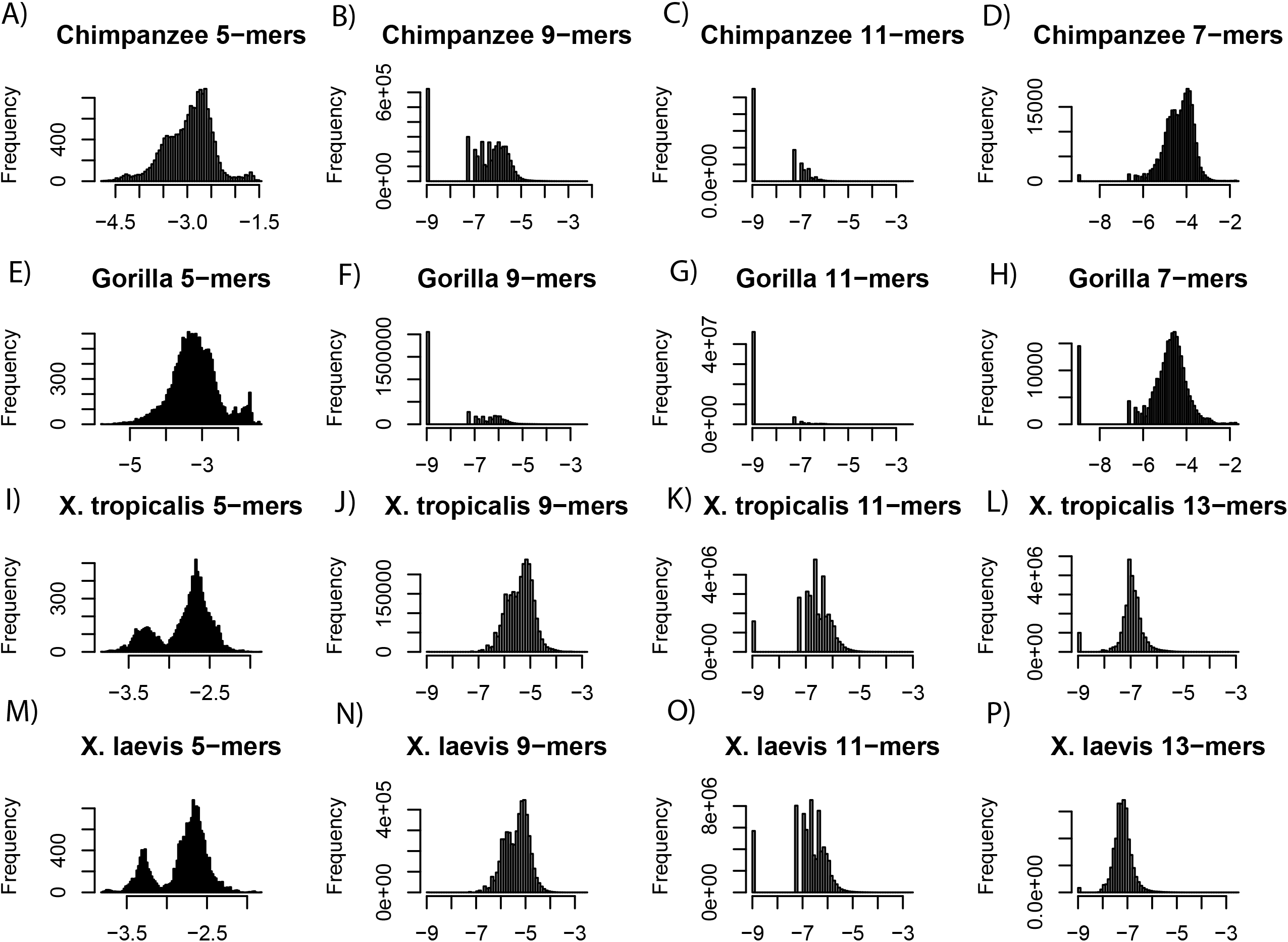
K-mer distributions in the subterminal sections of chromosomes Each panel shows the normalized k-mer density histogram for the species and k-mer length in the header. Each row shows the distribution for a species: a)-d) are Chimpanzee, e)-h) are Gorilla, i)-l) are *X. tropicalis*, and m)-p) are *X. laevis*. The first 3 columns are for k=5, k=9, and k=11 in each species. The 4th column shows the k-mer used for that species, 7-mers for primates and 13-mers for amphibians.

**Supplemental Figure 2.**
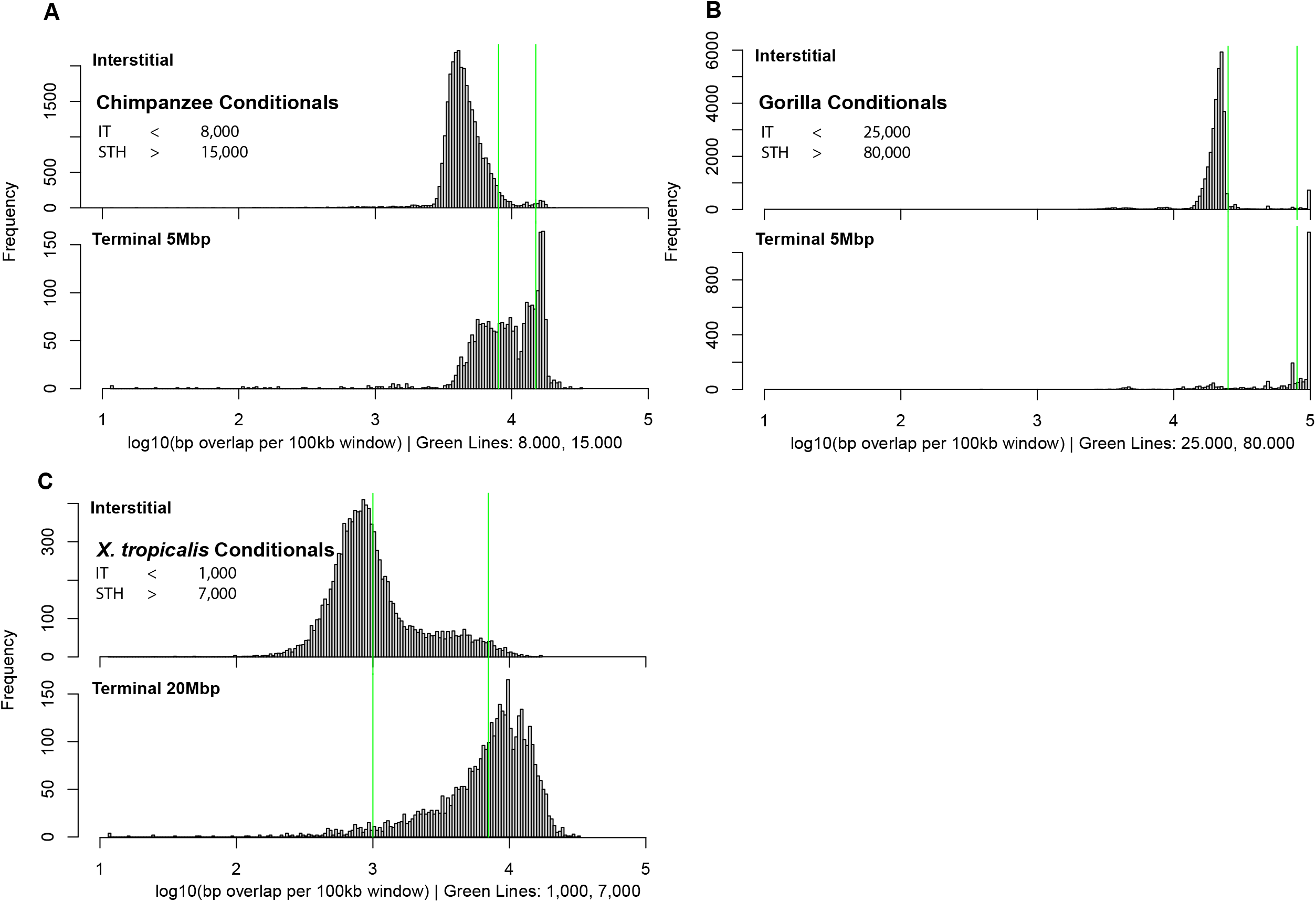
Diploid Hidden Markov Model conditionals Panels A, B and C show the number of bp of overlapped significant STH k-mers in each 100kbp bin within the interstitial (top) and the estimated STH search space (bottom) within Chimpanzee, Gorilla and *X. tropicalis*, respectively. Conditionals which determine the emission for the HMM are shown within each panel. Any 100kbp bins that are FALSE for any condition (falling between the green lines) will emit an Unknown signal, which will later be assigned by the HMM.

**Supplemental Figure 3:**
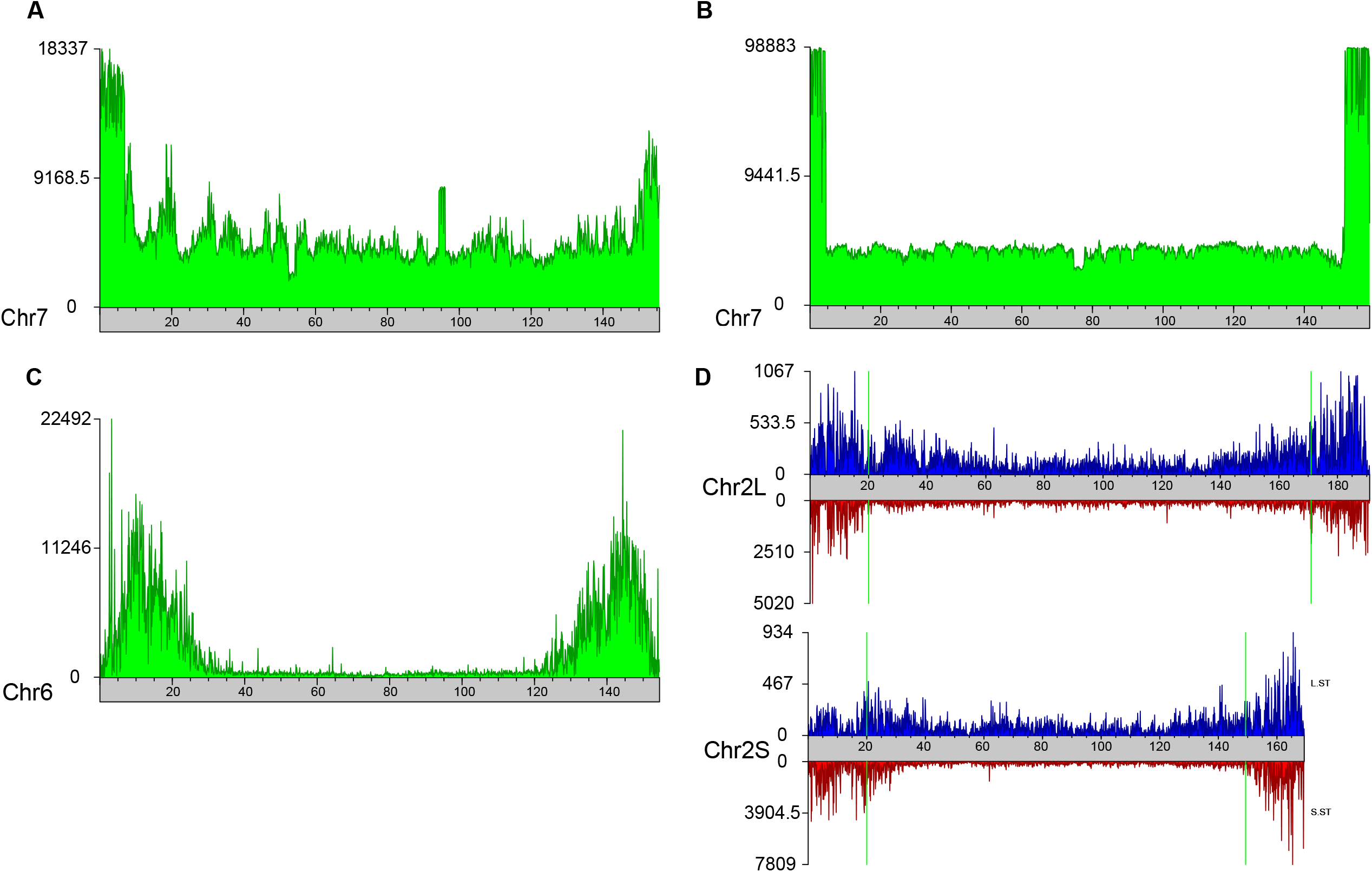
Training chromosomes A) Karyogram showing the density of subterminal 7-mers on chimpanzee Chr7, the training chromosome for the chimpanzee HMM. Each bin is 100-kbp. B) Karyogram showing the density of subterminal 7-mers on gorilla Chr8, the training chromosome for the gorilla HMM. Each bin is 100-kbp. For gorilla, the maximum subterminal density is 98,883bp/100,000bp bin. To show the full distribution, the gorilla y-axis is non-linear in scale. C) Karyogram showing the density of subterminal 13-mers on *X. tropicalis* Chr6, the training chromosome for the *X. tropicalis* HMM. Each bin is 100-kbp. D) Karyograms showing the density of L and S subterminal 13-mers on the two training chromosomes, Chr2L on top, Chr2S on bottom. L subterminal 13-mer density is shown on the top of the karyograms in blue, while S subterminal 13-mer density is shown on the bottom of the karyograms in red. Two chromosomes are necessary for *X. laevis* because the two subgenomes emit different k-mer signals.

**Supplemental Figure 4:**
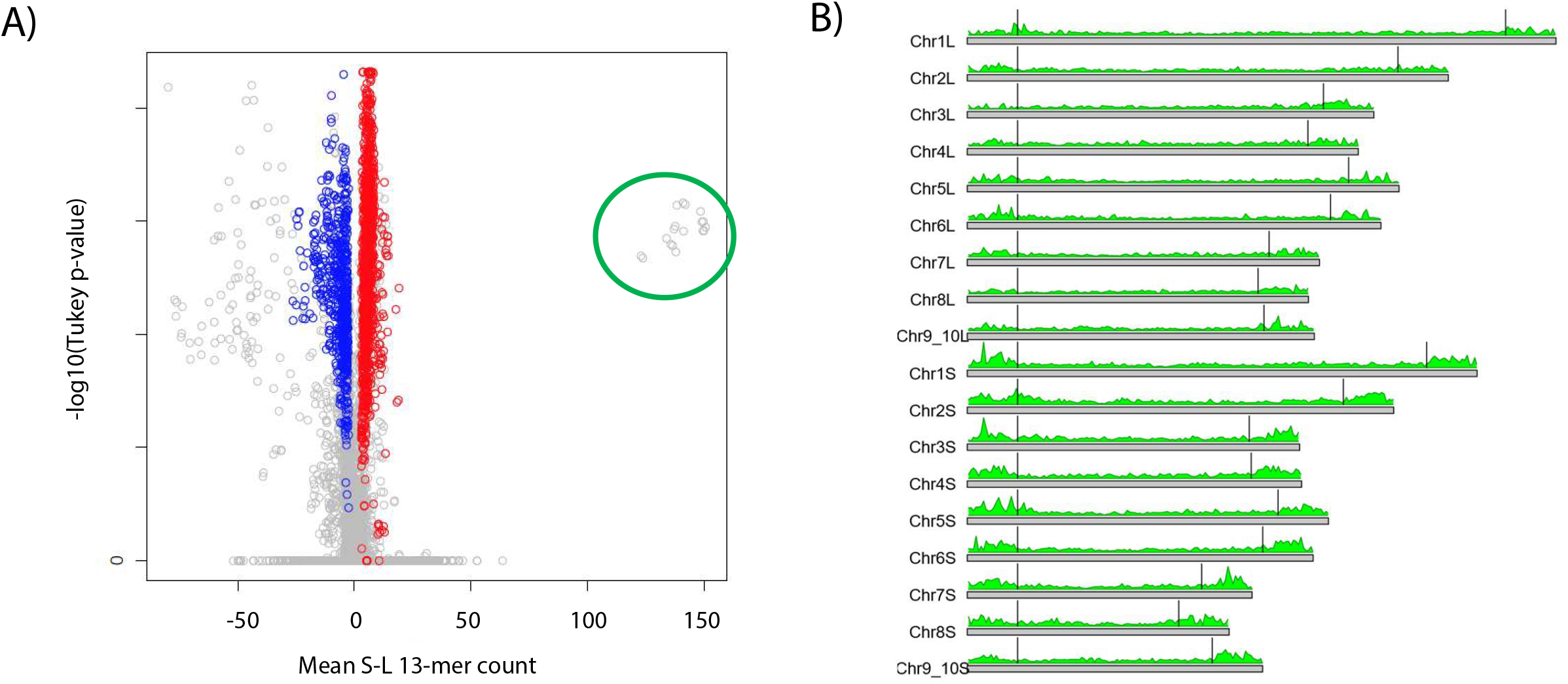
*X. laevis* subterminal heterochromatin size identification based on subgenome analysis outliers a) Volcano plot showing mean S-L 13-mer on the x-axis, and –log10(p-value) on the y using v10-2 *X*. laevis genome. P-values are calculated from Tukey’s HSD test following ANOVA testing for each 13-mers’ ability to differentiate the L and S subgenomes. Red and blue points are S 13-mers and L 13-mers that were significant in the previous genome build respectively. The green circle indicates outliers compared to previous analysis. b) Karyogram showing the density of 83 13-mers in the green circle from Supplemental Figure 4a, which build up in the terminal ends of the chromosome, potentially marking subtelomeres. Vertical lines are drawn at 20Mbp from the tips of chromosomes to show the initial range used to search for the subtelomeric sequences specifically.

**Supplemental Figure 5:**
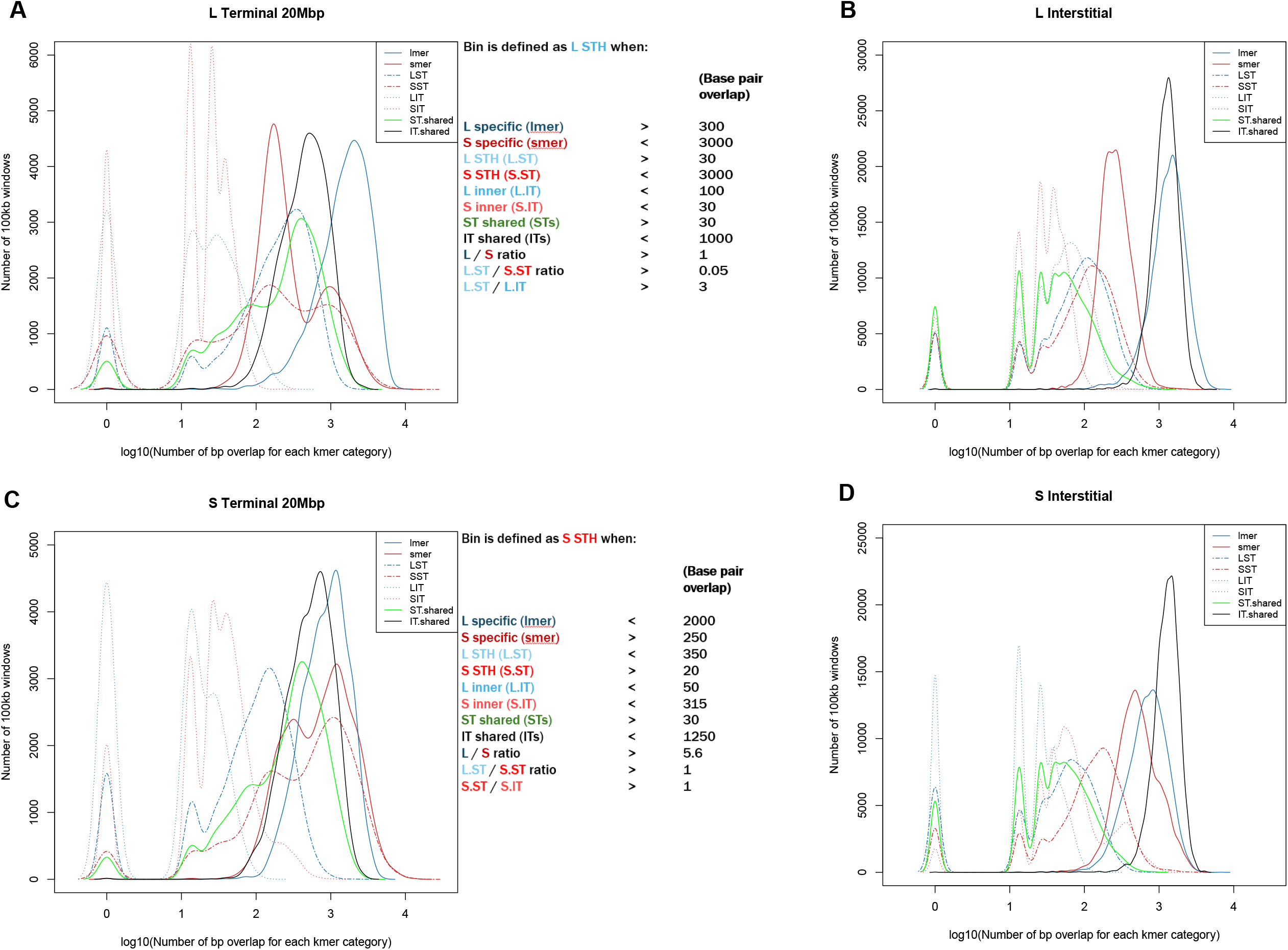
*X. laevis* HMM conditionals A) Kernel density plots (left) and HMM conditionals (right) for *X. laevis* terminal 20Mbp regions in the L subgenome. Each line is the kernel density plot for the labeled 13-mer group in the terminal 20Mbp of *X. laevis* L chromosomes, using the density in 100-kbp bins. The conditionals listed are the required values for a single 100-kbp bin to emit a “L-subterminal(LST)” signal. B) Kernel density plots (left) and HMM conditionals (right) for *X. laevis* terminal 20Mbp regions in the S subgenome. Each line is the kernel density plot for the labeled 13-mer group in the terminal 20Mbp of *X. laevis* S chromosomes, using the density in 100-kbp bins. The conditionals listed are the required values for a single 100-kbp bin to emit a “S-subterminal(SST)” signal. C) Kernel density plots for the interstitial segments of *X. laevis* chromosomes in the L subgenome. Each line is the kernel density plot for the labeled 13-mer group in the interstitial (middle) segment of *X. laevis* L chromosomes, using the density in 100-kbp bins. D) Kernel density plots for the interstitial segments of *X. laevis* chromosomes in the S subgenome. Each line is the kernel density plot for the labeled 13-mer group in the interstitial (middle) segment of *X. laevis* S chromosomes, using the density in 100-kbp bins.

