## Supplemental Notes for "Differential k-mer distributions enable identification of tetrapod subterminal heterochromatin"

**Supplementary Note 1: Choice of k in searching for subterminal heterochromatin**

The length of k in this analysis impacts the ability of the algorithm to recognize longer repeats. Generally, shorter lengths of k cause all k-mers to become common, and risks any subtle difference in density to become too small to recognize next to the high counts of the short k-mers. Longer lengths of k are generally more informative, at the cost of computational efficiency. The total number of possible k-mers, taking into account reverse complement sequences, is $2*(({1/4)}^{k})$ . Using this equation, k=13 requires 33,554,432 calculations per analysis step unless low count uninformative k-mers are first removed, even then typically leaving >1,000,000 k-mers to analyze per step. We have used k=13 and k=15 previously for subgenome identification across the eukaryotic tree, which always produced unimodal Gaussian distributions when looking at normalized counts for each chromosome ^1^, however k>=13 is not valid for analyzing lengths of sequence as small as primate subterminal heterochromatin due to most 13-mers not appearing in 5 million base pairs on average. We chose k=7 for these because this left few 0 count k-mers per terminal end. We investigated k=5, k=9, and k=11 and found all to be either bimodal or having too many 0’s in their distribution in the genomes analyzed here (Supplemental Figure 1). Excess 0’s in the subterminal distributions begin at k=9. The non-Gaussian k-distributions require non-parametric tests for significance, which lowers overall statistical power. This lower statistical power is sufficient for identifying the known pCht satellite in both gorilla and chimpanzee genomes however, speaking to the robustness of our algorithm.

**Supplementary Note 2: Primate Telomere-to-Telomere assembly subterminal heterochromatin analysis**

**Gorilla**

We obtained the Telomere-to-Telomere (T2T) genome from NCBI (NHGRI_mGorGor1-v2.0_pri; Supplemental Table 1). We defined the terminal 5Mbp of each non-acrocentric chromosome as the search space for the subterminal chromatin as this is the average size estimated in primates^2,3^. We split the chromosomes into 5p, 3p terminal 5Mbp and interstitial using blastdbcmd^4^. We counted 7-mers Jellyfish^5^, and normalized (k-mer/bp) the counts using R. The normalized counts produce bimodal distributions (Supplemental Figure 1). Using a non-parametric Kruskal-Wallis test^6^ followed by the post-hoc Dunn’s test^7^, we identified 496 subterminally enriched 7-mers, after applying the Bonferroni correction. We found their locations along chromosomes using a modified python script^8^. We merge overlapping k-mers via Bedtools^9^ merge (v2.30.0), as multiple k-mers may represent a single long repeat. We intersect this repeat bed file (via bedtools intersect) with a bed file that separates the genome into 100 kbp windows (created using awk^10^ and bedtools makewindows). We grouped resulting bed files into 100kbp bins using bedtools groupby. We made visualizations showing density along chromosomes using the kpArea() function from the karyoploteR package in R^11^. We define conditionals determining emission of each 100kbp bin based on the unique distributions in the terminal 5Mbp and interstitial regions (Supplemental Figure 2). We chose chromosome 7 as the training for the HMM which creates the emission table (Supplemental Figure 3). HMM probability tables are shown below. We compared to pCht as defined in Yoo et al.^2,3^ using blastn^4^ creating 3 groups: strict 6 or more matches to pCht (196/496), permissive 4 or more matches(434/496), and non-pCht with no matches (62/496).

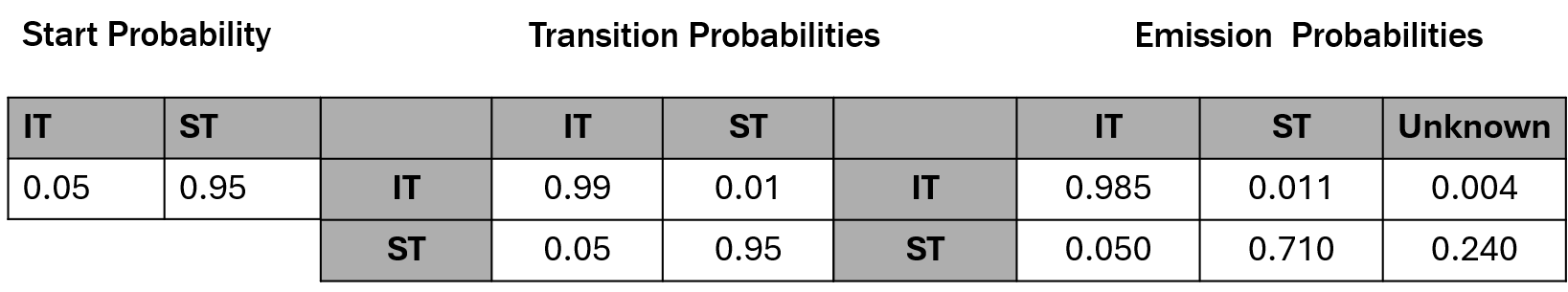

**Gorilla HMM probability tables**

**(left)** The table shows the start probabilities assigned for the HMM. This indicates the probability of assigning the first 100kbp window as subterminal heterochromatin (ST) or interstitial (IT).

**(middle)** The transition probabilities represent the likelihood that the HMM will transition from the row state into the column state given the assignment of the previous 100kbp bin.

**(right)** The emission probabilities are obtained from the training chromosomes (Supplemental Fig 3). The emission states are defined by the conditionals assigned to each specific species given their distributions (Supplemental Fig 2).

In order to assess if we could identify pCht without the existing data sets, we ran Tandem Repeat Finder^12^ (TRF) on the gorilla T2T genome using default parameters. We compared our subterminal k-mers to the TRF satellite consensi using BLASTN using the same parameters described above for pCht. Our subterminal k-mers pull down many TRF consensi, including multiple instances of pCht, however counting specific alleles is difficult due to TRF not enforcing a singular start position to its satellites. Because of this, we cannot estimate how many extra alleles of pCht were identified, or exactly how many extra satellites were identified using our method. However, we can report that 479 satellite consensi identified by TRF that contain our subterminal 7-mers have a significant BLAST alignment (evalue < 1e-2) to the pCht consensus while our subterminal 7-mers align to a total of 29,822 satellites, indicating many lower density subterminal satellites that are not related in sequence to pCht.

**Chimpanzee**

We obtained the T2T genome from NCBI (NHGRI_mPanTro3-v2.0_pri, Supplemental Table 1). We defined the terminal 5Mbp of each non-acrocentric chromosome as the search space as this is the average size estimated in primates^2,3^. We split the chromosomes into 5p, 3p terminal 5Mbp and interstitial using blastdbcm. We counted 7-mers using Jellyfish^5^, and normalized the counts using R. Normalized counts produce bimodal distributions. Using a non-parametric Kruskal-Wallis^6^ test followed by the post-hoc Dunn’s test^7^, we identified 207 subterminally enriched 7-mers, after applying the Bonferroni correction. We found their locations along chromosomes using a modified python script. We merge overlapping k-mers via Bedtools^9^ merge (v2.30.0), as multiple k-mers may represent a single long repeat. We intersect this repeat bed file (via bedtools intersect) with a bed file that separates the genome into 100 kbp windows (created using awk^10^ and bedtools makewindows). We grouped resulting bed files into 100kp bins using bedtools groupby. We made visualizations showing density along chromosomes using the kpArea() function from the karyoploteR package in R^11^. We define conditionals determining emission of each 100kbp bin based on the unique distributions in the terminal 5Mbp and interstitial regions (Supplemental Figure 2). We chose chromosome 7 as the training for the HMM which creates the emission table (Supplemental Figure 3). HMM probability tables are shown below. We comparison to pCht as defined in Yoo^3^ using blastn creating 3 groups: strict 6 or more matches to pCht (31/207), permissive 4 or more matches(140/207), and non-pCht with no matches (67/207).

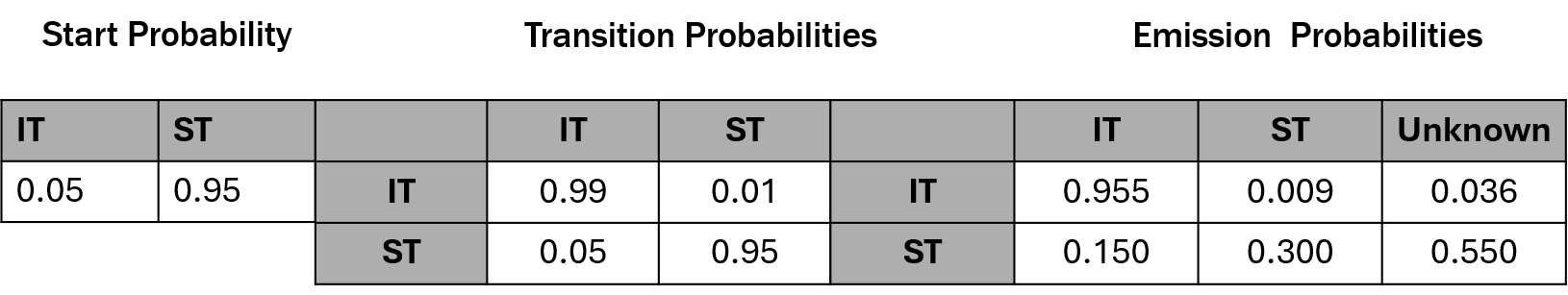

**Chimpanzee HMM probability tables**

**(left)** The table shows the start probabilities assigned for the HMM. This indicates the probability of assigning the first 100kbp window as subterminal heterochromatin (ST) or interstitial (IT).

**(middle)** The transition probabilities represent the likelihood that the HMM will transition from the row state into the column state given the assignment of the previous 100kbp bin.

**(right)** The emission probabilities are obtained from the training chromosomes (Supplemental Fig 3). The emission states are defined by the conditionals assigned to each specific species given their distributions (Supplemental Fig 2).

In order to assess if we could identify pCht without the existing data sets, we ran Tandem Repeat Finder^12^ (TRF) on the chimpanzee T2T genome using default parameters. We compared our subterminal k-mers to the TRF satellite consensi using BLASTN using the same parameters described above for pCht. Our subterminal k-mers pull down many TRF consensi, including multiple instances of pCht, however counting specific alleles is difficult due to TRF not enforcing a singular start to its satellites. Because of this, we cannot estimate how many extra alleles of pCht were identified, or exactly how many extra satellites were identified using our method. However, we can report that 989 satellite consensi identified by TRF that contain our subterminal 7-mers have a significant BLAST alignment (evalue < 1e-2) to the pCht consensus while our subterminal 7-mers align to a total of 30,014 satellites, indicating many lower density subterminal satellites that are not related in sequence to pCht.

**Supplementary Note 3: *X. tropicalis* subterminal heterochromatin analysis**

The genome we obtained from Xenbase (*X. tropicalis* v10.0 inbred) utilizes shotgun and long-read sequencing. The predicted STH regions are on average ~20Mbp according to a previous method based on segmental duplications^13^ so we use 20Mbp as our subterminal heterochromatin search space. We split chromosomes into 5p, 3p terminal 20Mbp, and interstitial using blastdbcmd. We count 13-mers using Jellyfish. We remove any k-mer with a total count of <100 as it is unlikely to be informative to our analysis and reduces computational processing time (Supplemental Note 1). We normalized and tested 2,760,255 13-mers via the ANOVA + TukeyHSD tests for their ability to partition between the terminal 20Mbp and the interstitial region, applying the Bonferroni correction. We found locations along chromosomes using a modified python script. We merge overlapping k-mers via Bedtools^9^ merge (v2.30.0), as multiple k-mers may represent a single long repeat. We intersect this repeat bed file (via bedtools intersect) with a bed file that separates the genome into 100 kbp windows (created using awk^10^ and bedtools makewindows). We group the resulting bed files into 100kp bins using bedtools groupby. We made visualizations showing repeat density along chromosomes using the kpArea() function from the karyoploteR package in R^11^. We defined conditionals determining emission of each 100kbp bin based on the unique distributions in the terminal 20Mbp and interstitial regions. We apply the HMM similarly to the primates and we calculate the emission probabilities using Chromosome 6 as the training chromosome (Supplemental Figure 3). We made comparisons to the 52bp satellite highlighted in Bredeson *et al.**^13^* similarly to the primate pCht satellite, using blastn to match 12 or more bases from each significant kmer. Relaxed criteria covers the entire 52-mer.

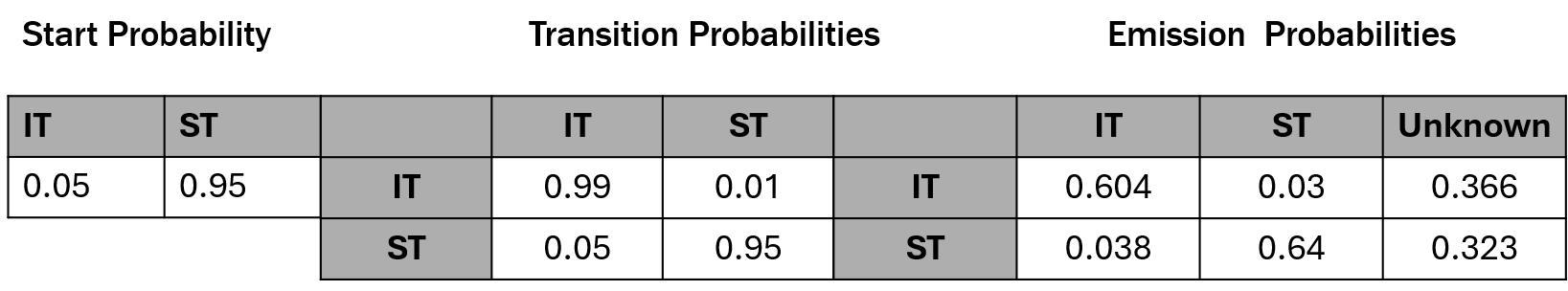

***X. tropicalis* HMM probability tables**

**(left)** The table shows the start probabilities assigned for the HMM. This indicates the probability of assigning the first 100kbp window as subterminal heterochromatin (ST) or interstitial (IT).

**(middle)** The transition probabilities represent the likelihood that the HMM will transition from the row state into the column state given the assignment of the previous 100kbp bin.

**(right)** The emission probabilities are obtained from the training chromosomes (Supplemental Fig 3). The emission states are defined by the conditionals assigned to each specific species given their distributions (Supplemental Fig 2).

In order to assess if we could identify the previously characterized 52-mer without the existing data sets, we ran Tandem Repeat Finder12 (TRF) on the *X. tropicalis* genome using default parameters. We compared our subterminal k-mers to the TRF satellite consensi using BLASTN using the same parameters described above for comparison to the Bredeson et al.^13^ 52-mer. Our subterminal k-mers pull down many TRF consensi, including multiple instances related to the 52-mer, however counting specific alleles is difficult due to TRF not enforcing a singular start to its satellites. Because of this, we cannot estimate how many extra alleles of the 52-mer were identified, or exactly how many extra satellites were identified using our method. However, we can report that 2,009 satellite consensi identified by TRF that contain our subterminal 13-mers have a significant BLAST alignment (evalue < 1e-2) to the 52-mer consensus while our subterminal 13-mers align to a total of 62,470 satellites, indicating many subterminal satellites that are not related in sequence to the previously characterized 52-mer.

**Supplementary Note 4: *X. laevis* subterminal heterochromatin analysis**

The genome we obtained from Xenbase (*X. laevis* v10.1 J-Strain) is assembled using a combination of whole genome shotgun sequencing and long-read sequencing via PacBio. In *X. laevis* the STH are estimated to be around 20Mbp based on similarity to X. tropicalis and preliminary data (Supplemental Figure 4), so we use 20Mbp as our search space for the subterminal heterochromatin. After identifying k-mers enriched in the STH regions, we use a Hidden Markov Model (HMM) to assign each region of the genome to being one of fours states: L-STH (LST), S-STH (SST), L-interstitial (LIT), or S-interstitial (SIT)^14^. We tried both 100,000 bp (100kbp) and 1,000,000 bp (1Mbp) bins for this analysis and found 100kbp to have a clearer visualization of the distribution.

We split chromosomes into 5p, 3p terminal 20Mbp, and interstitial using blastdbcmd. Using Jellyfish^5^(v2.3.1) we computed 13-mers in each region, whose normalized counts produced unimodal distributions . We remove any k-mers that occur <100 times as they are likely uninformative to our analysis. We normalized and tested 6,966,061 13-mers via the ANOVA + TukeyHSD tests for their ability to partition between the terminal 20Mbp and the interstitial region with respect to subgenomes L and S, applying the Bonferroni correction (Figure 1b). We found locations along chromosomes using a modified python script. We merge overlapping k-mers via Bedtools^9^ merge (v2.30.0), as multiple k-mers may represent a single long repeat. We intersect this repeat bed file (via bedtools intersect) with a bed file that separates the genome into 100 kbp windows (created using awk^10^ and bedtools makewindows). We group the resulting bed files into 100kp bins using bedtools groupby. Supplemental Figure 5 shows the kernel density plots for each k-mer group in the terminal 20MB of L and S chromosomes. Each of the regions described in Supplemental Figure 5, (including L and S interstitial), have a distinct distribution for each k-mer group. Thus, we use these distributions to differentiate between the interstitial and STH regions. We made visualizations showing repeat density along chromosomes using the kpArea() function from the karyoploteR package in R^11^. We defined sets of conditionals that can partition each region (interstitial and STH) with respect to subgenome (Supplemental Figure 5). K-mers partitioning L and S subgenomes from previous analysis^15^ are also included to better partition between the subgenomes. The HMM uses the sub-chromosomal k-mer densities discussed in Supplemental Figure 5 assigning 100kbp windows of the training set to either region (emission), in order to assign an observable process (path), in this case: finely mapping each individual 100kbp bins state (excluding the last 3’ bin which does not equal to 100 kbp) as either L or S STH (LST, SST) or L or S interstitial(LIT, SIT). The HMM uses a set of assigned probabilities (below) and a training data set which best reflects the expected chromosomal regions. In the case of *X. laevis*, chromosomes 2L and 2S show the clearest distribution of significant k-mers partitioning the terminal 20Mbp from the interstitial region (Supplemental Figure 4D), and thus, we choose them as the training set. The emission probabilities come from the training data. Our start probabilities favor STH, however, allow for the final bins to be interstitial. Our transition probabilities favor the same state the highest, then within subgenome transition, then cross subgenome transitions are least likely but still possible. Data was analyzed and Viterbi path calculated using HMM package in R^16^. HMM calls are visualized using the karyoploteR package in R^11^.

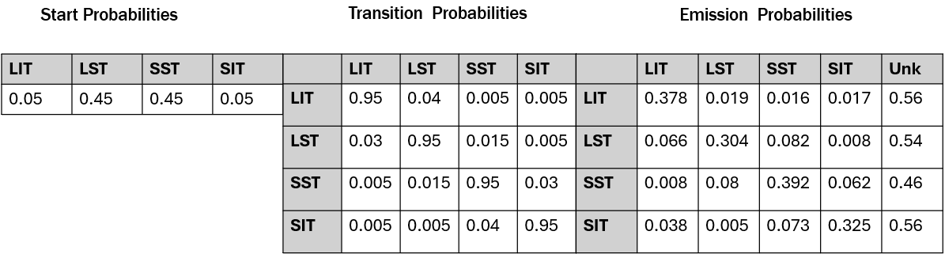

***X. laevis* HMM probability tables**

**(left)** The table shows the start probabilities assigned for the HMM. This indicates the probability of assigning the first 100kbp window as L subterminal (LST) or interstitial (LIT), S subterminal (SST) or interstitial (SIT).

**(middle)** The transition probabilities represent the likelihood that the HMM will transition from the row state into the column state given the assignment of the previous 100kbp bin.

**(right)** The emission probabilities are obtained from the training chromosomes (Supplemental Fig 3). The emission states are defined by the conditionals assigned to each specific species given their distributions (Supplemental Fig 2).

**7.**	**Dunn, O. J. Multiple Comparisons using rank sums. *Technometrics* 6, 241–252.**

**8.**	**dariober. bioinformatics-cafe/fastaRegexFinder at master · dariober/bioinformatics-cafe. *GitHub* https://github.com/dariober/bioinformatics-cafe/tree/master/fastaRegexFinder.**

**16.**	**Himmelmann, S. S.-D. L. HMM: Hidden Markov Models. (2022).**
